# Protons in gating the K_v_1,2 channel: a calculated conformational change in response to addition of a proton, and a proposed path from voltage sensing domain to gate

**DOI:** 10.1101/2022.03.31.486624

**Authors:** Alisher M. Kariev, Michael E Green

## Abstract

We have in the past proposed that protons constitute the gating current in the potassium channel K_v_1.2. Here we present a quantum calculation of a protonation change in a 311 atom section of intracellular S4-S5 linker, together with part of the T1 intracellular moiety of the channel. This proton shift leads to a hinge rotation in the linker, which in turn produces a separation of two amino acids, K312 and R326 (using the numbering of the 3Lut pdb structure). Two complete proton wires can then be proposed that would fully account for the gating mechanism with protons; the proton wires have as yet not been completely calculated. However, the path seems reasonably evident, based on the amino acids in the S4-S5 linker, which connects to the pore transmembrane S6 segment as well, and the T1 moiety of the channel, which is part of one proton path. This therefore also accounts for the T1 effect on gating. We had earlier shown how a proton could be generated from the VSD. Taken together the paths from the VSD to the gate show how the VSD can couple to the gating mechanism by having protons move between the VSD and the gate, closing the channel by both producing the hinge rotation and providing electrostatic repulsion to an incoming K^+^ ion. The protons move under the influence of membrane polarization/depolarization. Taken together, this makes our previous model much more detailed, specifying the role of particular amino acids.

## Introduction

Voltage gated potassium channels of the type we are discussing have four voltage sensing domains (VSD), each with four transmembrane (TM) segments; one of these, labeled S4, contains several arginine amino acids that are presumably positively charged, while there are counter charges on other TM segments, forming salt bridges[1]. A transient capacitative current accompanies gating of these and of sodium channels [2, 3], so there is no question that charges are moving, and there is good reason to believe these to be positive charges. A conventional model has become more or less generally accepted [4] in which the channel opens by having the S4 segments move in the extracellular direction in response to depolarization of the cell membrane within which the channel sits. To close the channel, the S4 segments move in the opposite direction in response to repolarization of the membrane. We have suggested that caution is required in interpreting the evidence for this model [5], and have suggested an alternative method for gating [6, 7] in which protons provide the mobile positive charges, rather than S4 segments as a whole. We suggest that what is moving are at least two, and probably three, protons, for each of the four domains. Based on quantum calculations, it is possible to show that at least one proton can be generated from an arginine-glutamate-tyrosine triad of amino acids in the VSD [7, 8]. It is known that the H_v_1 channel, which is very similar to the K_v_1.2 VSD in its upper half (approximately) does in fact transmit protons, as does bacteriorhodopsin. The H_v_1 results, albeit with not quite the same interpretation, have been reviewed by deCoursey [9]. All three (K,1.2, H_v_1, bacteriorhodopsin) have similar structures for their proton transport sections, a point that will be considered later. Helms and Gu found a double proton wire in green fluorescent protein, a case that is an even closer analogy to the proposal presented here [10] A number of examples of triads of amino acids have been proposed; Chanda and coworkers have explicitly worked out the thermodynamics of an arginine, glutamate, tyrosine triad in the context of gating [11]. This is the same triad that we found produces a proton by way of tyrosine ionization [7, 8]. In fact, proton paths in multiple proteins are known, far too many to cite here; however, we can cite at least some relevant recent work [12] [13–16]. In addition there are cases of proton transport producing allostery, with, as examples, influenza M2 virus [17], the NHE1 and NHE3 Na^+^/H^+^ exchanger transporters [18, 19] and other cases [20, 21]. Our proposed mechanism, although it differs from standard channel models, does not involve any new physical or biological principles [22]. Proton paths can be found both in H_v_1 and in K_v_1 for the sections where the two diverge. In H_v_1 the path proceeds fairly directly to the membrane headgroup region, where the proton can be released [23], while there is a bend in the path in K_v_1. It is possible to find what appears to be a proton wire in the intracellular portion of the K_v_1 channel that continues from the VSD to the gate, and it this path that we discuss here. Our proposed path includes several amino acids that are known to be important in gating, from experimental mutations [22, 24] Proton binding has been found in other channel systems, like the just cited bacteriorhodopsin, known to transmit protons [25, 26]. This is accompanied by proton transport [27], which can be unidirectional [28]. What is interesting is that triads of amino acids, often a salt bridge plus tyrosine, occur in several of these cases; it appears that triads are able to transmit protons effectively. In this work, we see a path composed not only of triads, but in significant part of longer amino acid sequences, with four or even five, amino acids, that could function as part of a proton wire. Adding one, or sometimes two, water molecules to bridge these groups, makes the paths complete. Two residues can be identified as necessary termini for one proton chain, as we will see in the results. In addition, K_v_1.2 has, at the gate, a group of two asparagines and a serine. The three oxygens in the side chains look as though they would necessarily play a role in proton transport. This section has not yet been calculated; we note their presence at this point, but will include them in our discussion.

## Methods

We have carried out quantum calculations on a critical 311 atom section of this path, and can easily see what appears to be a proton wire for the remainder; in fact, two paths can be found, one of which involves the T1 moiety of the intracellular section of the channel. It has been known for many years that this segment is somehow involved in gating [29]. The quantum optimization (energy minimization) of this 311 atom portion of the intracellular section of the channel includes much of the linker of the VSD to the pore (the S4-S5 linker, together with a small part of transmembrane segment S6); the optimization was done at HF/6-31G** level, using Gaussian09 [30]. The calculation starts from the X-ray structure of the open conformation, with the protons added by a normal mode analysis in the 3Lut structure [31]; Our closed model places the protons on specific amino acids, E327 (by itself, still an open conformation) and the K312—R326 pair (see Fig. 2). The fully closed conformation may take one extra proton for the S411-N412-N414 group at the corner of the gate. There is evidence for the importance of the two specific residues that we consider to be protonated/deprotonated, so that the choice of proton locations is not arbitrary, but is in accordance with experimental evidence. The section that was calculated corresponds to one domain. The total charge on the 311 atom system as calculated is zero whether closed or open; in the calculation, charge is moved, but not added.

**Fig. 1.**
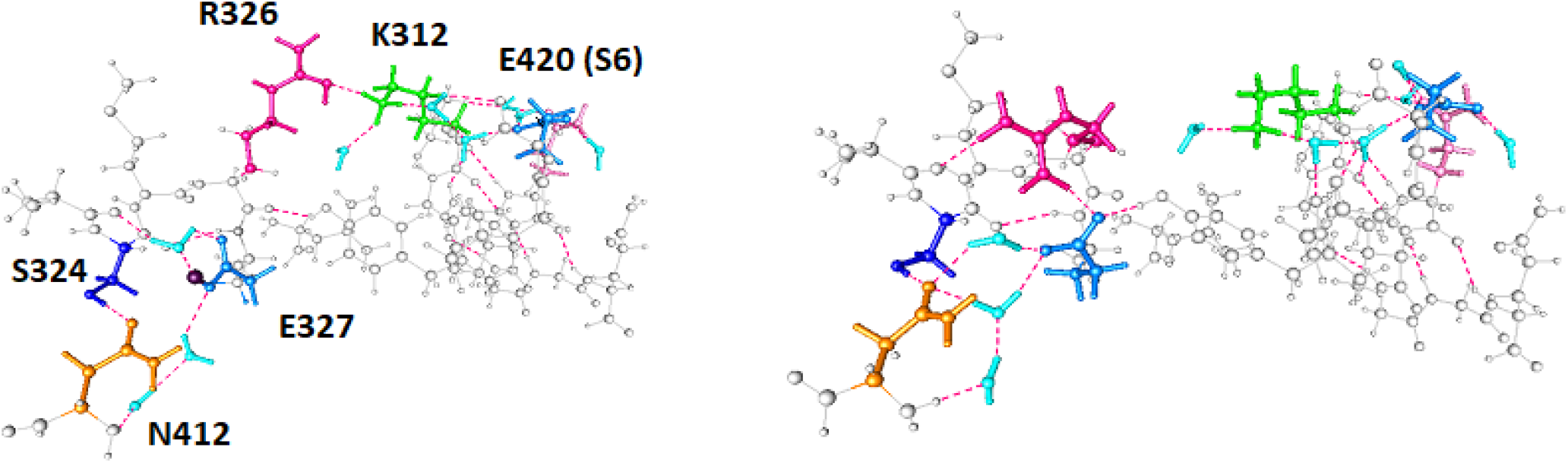
The effect of shifting a single proton: A) one proton on K312 and R326, plus one on E327; B) two protons on K312 and R326. Net charge is 0 on both A) and B). The corresponding amino acids have the same color in both. Note the separation between K312 and R326 in the doubly protonated figure B. This should be enough to matter in gating. R326 is clearly reoriented and acting as a hinge, and the N414 (not shown in this figure—see Fig. 2) extends beyond the position in the unprotonated state.

**Fig. 2:**
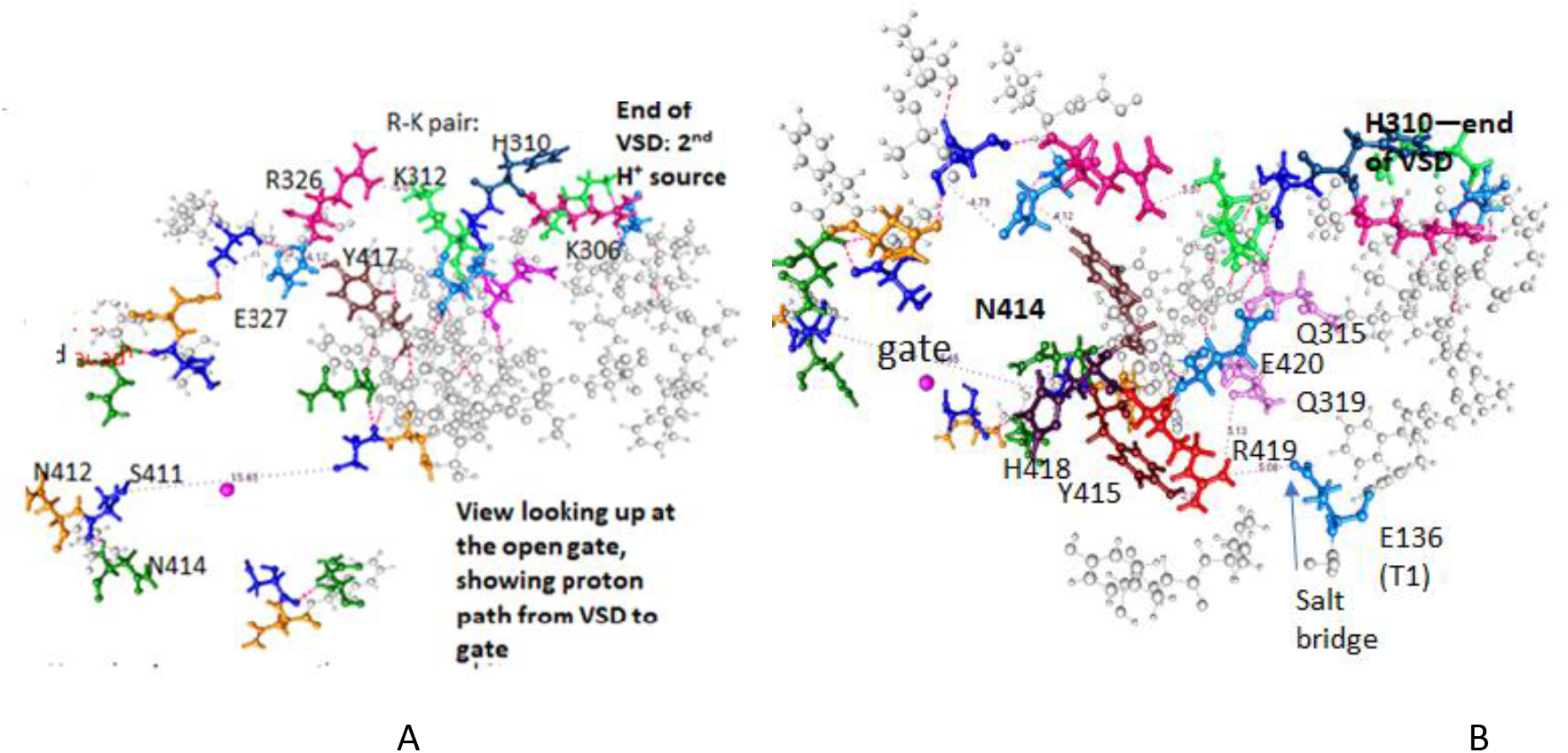
Two views of the X-ray structure (not energy minimized) of the entire proton paths from the end of the VSD to the gate: **A**, looking up from the intracellular side into the gate; **B**, the figure is rotated 90°^-^to give a side view. Both are the unprotonated open channel, and consistent with the 311 atom completed calculation.

## Result of the computation

One key feature is that the X-ray structure shows a lysine (K312) and an arginine (R326) in close proximity, with distance 5.07 Å for the K312 nitrogen to the nearest R226 nitrogen. If only one proton is shared by the two residues, the calculation shows the structure of the calculated energy minimum is fairly close to that of the X-ray structure, with the 5.07 Å N – N distance just mentioned calculated as 2.98 Å. If both are protonated, there is a large separation accompanying a hinge motion, separating the charges by 5.8 Å more, to 8.78 Å. The residues at the gate are on a relatively long lever arm, about 10 Å from the hinge to the end, so that the residues at the corner of the gate swing a glutamine (N412) down approximately 3 - 4 Å, slightly more if the hydrogens are included; the details are best seen in the figures, with this point clearest in Fig. 3. There is evidence that this should be sufficient to close the gate [32]. The second proton wire is kept constant in this calculation.

**Fig. 3:**
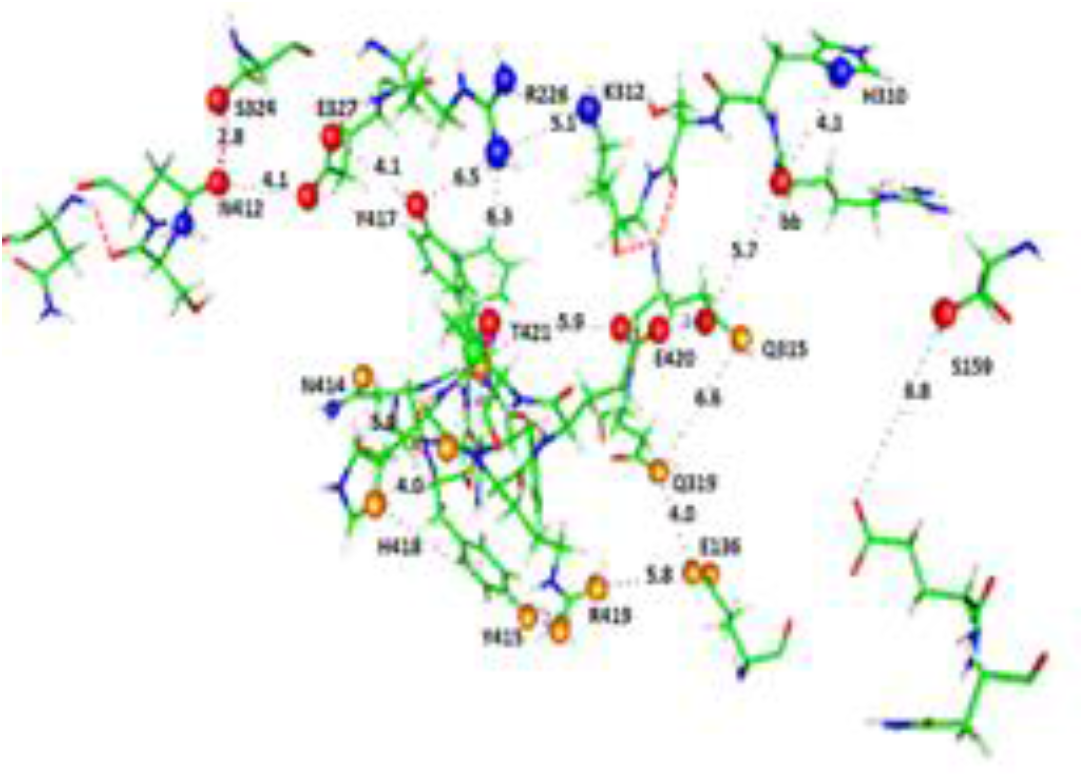
This shows the transitions in detail for the two paths, with key residues and distances, rounded to the nearest 0.1 Å, labeled. The distances are from the X-ray structure, not the optimization, but for the open state these agree fairly well. Note the key roles of H418 and E327; these have been identified experimentally by Lee et al as critical for gating[33]. However, all of the labeled residues are important in the paths; the R226 – K312 pair can be seen at the top; the N412 and N414 are probably the last step in transmitting the proton to the gate, if needed to close the channel.

One more proton is almost certainly necessary for gating, on the glutamate E327; the order in which the protons must be added to close the gate may be unimportant, but at this point it is not possible to say anything about the order in which the protons are added. A third proton may be added to the N412, N414, S411 group at the corner of the gate, (which has not yet been calculated), making it appreciably more positive. It is possible, without doing a quantum calculation at the center of the gate itself, which would require more atoms than it is feasible to include at the moment, to do a classical estimate, leading to several kBT at least of energy of interaction with a positive ion. However, the exact value depends on the local dielectric constant. It is known that confined water leads to a reduced dielectric constant, but the extent to which this affects the local potential requires an actual calculation. At present we allow for the possibility, even the likely presence, of a third proton, but must leave that for now with a qualitative argument. However, a role for two protons is seen in the section that has been calculated, providing a mechanism for going between the closed and open states. This allows the S4-S5 linker to play its key role without requiring it to fold in an improbable manner. It also accounts for the role of T1, which is known to be important, but is generally not accounted for in proposals for the VSD to gate link.

## Proposed complete proton wire from the VSD to the gate

Figure 2 shows views of the complete intracellular proton wires we suggest, including sections that remain to be calculated. At no point are more than two water molecules required to complete the gaps between the protein residues, and there is certainly water available at the intracellular surface of the membrane. While the parts of the wire that have not been calculated are, for this reason, at present hypotheses, they are consistent with everything we know, and consistent with the calculation that has been completed.

This gating model includes the effects of certain known mutations (H418 and E327 are of particular importance [33]) as well as the fact that the T1 segment is involved with gating, something not accounted for in essentially any version of the standard model. The standard models have some difficulty in showing just how the S4-S5 linker manages to transmit the force of an S4 motion to the gate, although it must do so in some manner. There have been proposals [34] for methods of mechanically having the linker press against the gate, closing it, but there appear to be alternative modes of motion for the linker; in a mechanical force model, the linker would have to be rigid, and there are gaps in the geometry that suggest that this would be difficult. For a proton path, water fills these gaps, and water can always be part of a proton wire.

At most two water molecules are needed to connect amino acids that can be part of the proton wire.

## NBO calculations

Natural Bond Orbital calculations[35] were carried out on the 311 atom optimized structures to determine atomic charges, using B3LYP/6-311G**. Table 1 summarizes the key results.

**Table 1.** Charges on key amino acid side chains

| Case* | Amino acid side chain | Atoms included | total charge |
| --- | --- | --- | --- |
| 1 | R226 | C(NH<sub>2</sub>)<sub>2</sub> | +0.853 |
| 2 | R226 | C((NH<sub>2</sub>)NH** | +0.071 |
| 1 | K312 | NH<sub>3</sub> | +0.599 |
| 2 | K312 | NH<sub>3</sub> | +0.621 |
| 1 | E327 | COO | -0.733 |
| 2 | E327 | COOH** | -0.012 |
| 1 | E420 | COO | -0.755 |
| 2 | E420 | COO | -0.756 |
| 1 | N315 | CONH<sub>2</sub> | +0.033 |
| 2 | N315 | CONH<sub>2</sub> | +0.028 |

\*Case 1: proton on (K312/R226) pair; Case 2: proton on E327
\*\*salt bridge

## Specific amino acid groupings that participate in the proton path

There are a number of groups of three, four, or five acidic or basic amino acids that compose much of the two proton paths. In addition to the acids and bases, there are amino acids that have hydrophilic side chains, serine and tyrosine, as well as asparagine. Actually, tyrosine can function as an acid, albeit weaker than glutamate and aspartate. Serine is important only in combination with two other amino acids that can act as acids or bases. It appears that the N412, N414, S411 group referred to earlier could act as a base and accept a proton, although, as noted, this remains to be calculated; the group could also act as part of chain to pass on a proton. Table 2 lists amino acid strings that are part of the paths from the VSD to the gate.

**Table 2.** Strings of Amino Acids in the Voltage Sensing Domain to Gate Pathways

| *String* | *Location* |
| --- | --- |
| E183 -- R297 -- Y266 -- E226 -- R303 | Within the VSD; produces a proton[8] |
| D259 -- R309 -- K306 -- E236 -- R240<br>- S159 -- Y155 | Intracellular end of VSD; upper path, and S159<br>branch point transition to lower main path |
| E327 -- Y417 -- R326 -- E420 | Upper path, near gate |
| Y415 -- R419 -- E136 | Lower path; includes E136 from T1 |

Table 1 does not show any great surprises: the charge goes with the proton. We can give a rough estimate of the repulsion between K312 and R226, with both protonated. With the distance taken to be an average over the atoms, we see that (depending on what the dielectric constant is estimated to be, but at that distance there is almost nothing between the two residues, so a very small dielectric constant is justified, perhaps as little as 2, but even if 4, we get a strong repulsion), there is of the order of 100 kBT energy, surely enough to force motion. At the end of the move, the distance at least doubled, and, more important, the dielectric constant increased. The increased distance permits water to fit between them, to allow these residues to be stable in their new positions. This water would not be so rigidly held, so the rotation would allow the water to be a better dielectric. A reasonable estimate suggests the repulsion should be about an order of magnitude less; it is not yet possible to achieve a really quantitative estimate.

Table 2 does not quite show the complete paths. In addition to up to two water molecules connecting these strings, the end of the path includes two residues, H418 and E327; only the latter is represented above. It is not hard to see how these are close enough to be the acceptor for each of the paths. E327 is within the upper path’s transfer range, while H418 could be reached via the lower path.

The transfer of a proton by just one amino acid is an advance along a path of 4 to 5 Å, not a negligible step in the context of the distances here. There are strings of acidic or basic amino acids that constitute parts of the paths (Table 2) although not directly connected to each other. Fig. 3 shows gaps principally of the size of one water molecule, counting the distance from one polar atom to another on the hydrophilic amino acids on either side of the water molecule; note how many are in the 5.8 – 6.8 Å range. There are one or two gaps that could accommodate two water molecules; possible folding makes these distances less well defined. There are apparently no gaps larger than this between amino acids that could be part of a proton wire. The relative paucity of hydrophobic amino acids in the linker suggests that a mechanical push from VSD to gate is not likely to be correct; hydrophobic amino acids would participate in a mechanical push at least as well as hydrophilic amino acids. With the gaps, it is likely that pushing on the linker could produce folding rather than pressure on the gate to the extent that it would move the backbone at all. Examining the positions of the amino acid strings, and adding water where it is most likely to be hydrogen bonded, produces two complete proton paths from VSD to gate. This is supported by an additional point: The hinge motion at the K312 – R326 junction shows how a conformational change in this system can be linked to proton transport and thence to gating. While this does not mean that electrostatic repulsion at the gate, or relatively rigid binding of water molecules near the gate, do not also contribute, it does mean that proton gating is compatible with a local conformational change near the gate that could have consequences down into the gate. The hinge motion itself is quite large in the calculations of open (one proton between K312 and R226, giving nitrogen to nitrogen distance 2.98 Å) and the proposed closed state (two protons, same atoms, distance 8.78 Å). The X-ray distance is 5.07 Å. A possible reason for the short distance in the calculation of the open structure is the absence of a nearby water molecule that would separate the two groups by an additional approximately 2 Å without being hydrogen bonded to them. It is clear in any case that the added proton can produce a hinge motion that appears large enough to close the gate, or at least make the major contribution to closing the gate. In addition, we see in the X-ray structure of the putatively open state the three oxygens of the carbonyl of the Q315 amide group and the two oxygens of E420, one of which is only 3.46 Å from the Q315 carbonyl oxygen, and the other just slightly further away. This could be accounted for by adding a third proton to the structure.

We now have reasonable termini for the two paths. The upper path does not need to extend to the gate, since the K312 – R266 pair already appears to produce the necessary effect, moving several residues several Angstroms when protonated. For the lower path, the two glutamines at the corner of the gate, N412 and N414, but especially the latter, transmit the proton to H418, which would account for the importance of this residue. A possible third proton would stop at E327, also known to be critical, but it is possible that this residue transmits the second proton to H418, rather than becoming protonated itself. Further computations are being done to answer this question.

## Conclusion

It is possible to find a gating mechanism for the K_v_1.2 channel that differs from the standard models, by which it is possible to interpret essentially all the evidence on gating, including that which has been adduced to support the standard models; by extension, it is likely to apply to other channels as well. In this alternative, the gating current consists of the motion of protons, while there is essentially no physical motion of the S4 segments of the VSD. The proton current, and proton gating, appear to solve certain difficulties with the standard S4 motion models. One such is the failure of standard models to consider involvement of the T1 intramolecular moiety of the channel. We have discussed the evaluation of evidence on gating in earlier work, including both evidence that has been adduced in support of the standard models, and evidence that supports the present model directly [5, 7, 8]. In other words, the proton model appears to be in accord with all experimental evidence, although alternate interpretations of the evidence, or at least some of it, are more commonly accepted. In addition, we have cited several cases, in bacteriorhodopsin, cytochrome c, and H_v_1, in which similar hydrophilic chains lead to water clusters that are part of proton paths, so that the entire mechanism is hardly unprecedented, except in the context of channel gating models. This model is supported by a quantum calculation for a key section of the paths, showing the most critical step.

## ACKNOWLEDGEMENT

Center for Functional Nanomaterials, which is a U.S. DOE Office of Science facility, and the Scientific Data and Computing Center, a component of the Computational Science Initiative, at the Brookhaven National Laboratory under contract no. DE-SC0012704 and resources at the High Performance Computation facility at the City University of New York

